# Loss of systemic anti-viral immunity and LMP1-driven suppressive myeloid tumour niches converge to shape the immunobiology of EBV^+^ diffuse large B-cell lymphoma

**DOI:** 10.64898/2026.01.26.701732

**Authors:** Éanna Fennell, Soi C. Law, Alexander C. Dowell, Matthew R. Pugh, Christina M. Enright, Aisling M. Ross, Aoife S. Hennessy, Ciara I. Leahy, Nadezhda Nikulina, Alex G. Richter, Lucia Mundo, Lorenzo Leoncini, Stefano Lazzi, Maria Chiara Siciliano, Stefan D. Dojcinov, Leticia Quintanilla-Martinez, Ken H. Young, Maher K Gandhi, Graham S. Taylor, Paul G. Murray

## Abstract

Epstein-Barr virus (EBV)-positive diffuse large B-cell lymphoma (EBV⁺DLBCL) is an aggressive lymphoma with poor outcomes and an incompletely understood pathogenesis, frequently attributed to immunosenescence. However, its occurrence across all age groups suggests alternative mechanisms. Here, we integrate functional profiling of peripheral antiviral T-cell immunity with high-dimensional spatial proteomics and mechanistic *in vitro* modelling to define the immunological landscape of EBV⁺DLBCL. We show that both EBV⁺ and EBV⁻DLBCL patients exhibit broad impairments in antiviral T-cell responses compared with healthy controls, affecting latent and lytic EBV antigens as well as non-EBV viral targets, with deficits most pronounced in EBV⁺ patients. Spatial proteomic analysis revealed that EBV⁺DLBCL harbours a profoundly immunosuppressive tumour microenvironment characterised by relative loss of intratumoural CD8⁺ T cells, expansion of PD-1⁺ regulatory and exhausted T-cell populations, and dense aggregates of PD-L1⁺/IDO1⁺ macrophages. Compared with EBV⁺ classical Hodgkin lymphoma and infectious mononucleosis, EBV⁺ DLBCL displayed the most marked macrophage-associated immunosuppressive signature and the lowest T-cell density. Suppressive myeloid niches were preferentially enriched around LMP1-expressing tumour cells, a feature not observed in the other EBV-associated conditions. Together, these findings indicate that EBV⁺DLBCL is driven by the convergence of systemic antiviral immune dysfunction and an LMP1-dependent suppressive tumour microenvironment.

## Introduction

EBV^+^ DLBCL is an aggressive B-cell malignancy characterised by expression of the EBV latent genes and a poor response to standard immunochemotherapy. EBV^+^ DLBCL was first described to occur exclusively in older adults, leading to the prevailing model that immunosenescence permits the outgrowth of EBV-transformed B-cells^1–5^. Supporting this, high circulating EBV DNA viral loads, potentially reflecting impaired immune control of EBV lytic replication, were found to be associated with worse patient outcomes^6^. However, EBV^+^ DLBCL is now recognised to occur across all age groups, including younger immunocompetent individuals, prompting its reclassification as EBV^+^ DLBCL, not otherwise specified (NOS)^7^. However, direct evidence describing the extent, specificity, and functional integrity of peripheral EBV-specific T-cell immunity in EBV^+^ DLBCL remains limited, leaving unresolved whether systemic immune deficiency is a significant feature of the disease.

In parallel, the TME of EBV^+^ DLBCL tumours was shown to exhibit transcriptional and phenotypic signatures of local immunosuppression ^8–10^. EBV is known to recruit a broad spectrum of immune cells, including T-cells and myeloid cells driven in part by EBV-induced alterations in chemokine expression^11–13^. *In vitro* studies show that several of these chemokines, such as CCL20 and CCL22, are regulated by the EBV-encoded latent membrane protein 1 (LMP1)^14^, a viral oncogene expressed by the tumour cells in almost all cases of EBV^+^ DLBCL^15^. Prior studies have also described increased checkpoint expression and M2-skewed macrophage infiltrates in EBV^+^ DLBCL. However, the spatial organisation and mechanistic relationships between tumour cells, T-cells and myeloid compartments have not been adequately resolved^12, 13, 16, 17^. Moreover, how the EBV^+^ DLBCL TME differs from that of EBV⁻ DLBCL or from other EBV-associated lymphoproliferative disorders is not known.

As a result, the field lacks a unified model explaining whether EBV^+^ DLBCL emerges from systemic immunological failure, from EBV-driven local immune evasion, or from the convergence of both processes^18^. To address these gaps in knowledge, we integrated detailed functional profiling of circulating T-cell immunity with high-dimensional spatial proteomic analysis of DLBCL tumours. We found that patients with DLBCL exhibit broad reductions in antiviral memory T-cell responses, spanning multiple EBV and non-EBV antigens, which was more pronounced in EBV^+^ DLBCL. This systemic deficit coexists with a profoundly immunosuppressive TME enriched for PD-1^+^ regulatory T-cells, PD-L1^+^/IDO1^+^ macrophages, and SPP1-expressing myeloid cells that cluster around LMP1-expressing tumour B-cells. Together, these data reveal that EBV^+^ DLBCL is characterised not by a single dominant failure but by the convergence of systemic antiviral T-cell impairment and a highly structured, LMP1-driven suppressive TME.

## Results

### DLBCL patients have impaired T-cell responses to EBV and non-EBV viral antigens, compared with healthy donors

To characterise systemic antiviral immunity in patients with DLBCL (pre-treatment and without human immunodeficiency virus (HIV) and immunosuppression; Supp. Table S1), we quantified peripheral T-cell responses to a panel of EBV and non-EBV viral antigens using IFN-γ ELISpot assays (Fig. 1; Supp. Table S3). Tumoural EBV status was determined by *in situ* hybridisation for the Epstein-Barr virus encoded RNAs (EBERs). We found that mitogenic responses to phytohemagglutinin (PHA) were comparable across healthy donors and in both EBV^-^ and EBV^+^ DLBCL patients, indicating intact global T-cell function. In contrast, antigen-specific profiling revealed broad impairments in antiviral immunity in the lymphoma patients. T-cell responses to the EBV latent nuclear antigens, EBNA1 and EBNA2, were significantly reduced in both DLBCL groups compared with healthy donors, and in EBV^+^ DLBCL compared with EBV- DLBCL. In contrast, T-cell responses to EBNA3A in EBV^+^ DLBCL were significantly reduced compared to EBV^-^ DLBCL and healthy controls. T-cell responses to the latent membrane proteins, LMP1 and LMP2 were also significantly reduced but only in patients with EBV^+^ DLBCL compared to healthy controls. Deficits were also pronounced for the EBV lytic antigens. Thus, recognition of the immediate-early protein BZLF1 was profoundly diminished in DLBCL patients and nearly absent in many EBV^+^ cases. Responses to the late lytic antigen gp350/450 were significantly lower in patients with EBV^+^ disease. However, impaired immunity extended beyond EBV-specific T-cell responses and included poor responsiveness to antigens from other herpesviruses (varicella-zoster virus and cytomegalovirus) as well as influenza. Analysis of T-cell responses in a second, independent cohort of healthy donors and DLBCL patients (also pre-treatment and without HIV and immunosuppression; Supp. Table S2) revealed a similar trend of reduced antiviral reactivity (Extended Data Fig. 1). Together, these findings demonstrate that DLBCL patients are characterised by a broad suppression of circulating antiviral T-cell responses that is especially pronounced in patients with EBV^+^ disease, and that may contribute to the impaired immune surveillance associated with tumour development. Importantly, these deficits were not explained by age, as responses were similarly reduced in younger patients, indicating that systemic immune dysfunction extends beyond classical immunosenescence.

**Figure 1.**
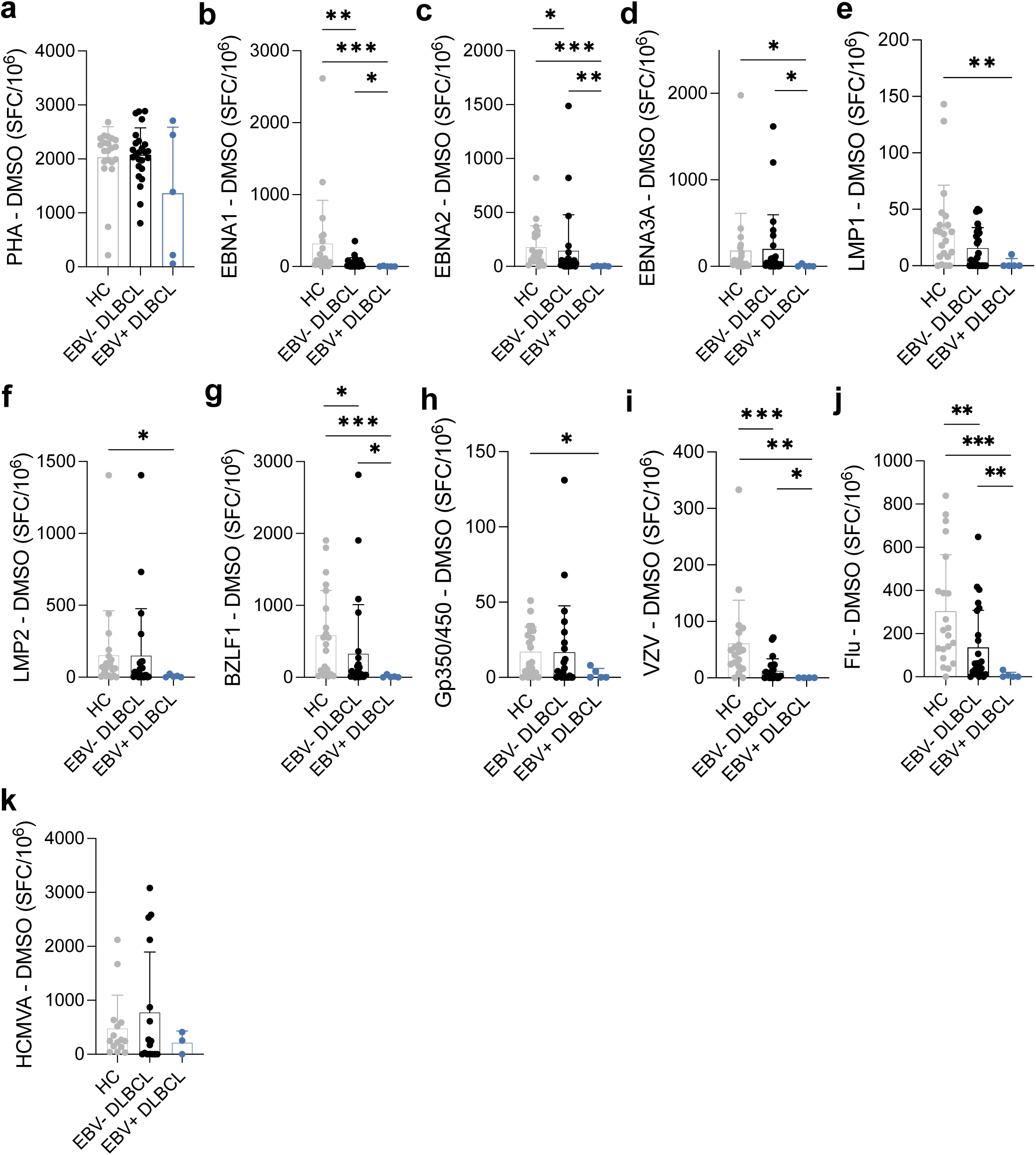
Frequency of T-cells specific for viral antigens in EBV^+^ and EBV^-^ DLBCL patients compared with healthy controls. **a - k.** *Ex vivo* IFNγ ELISpot response to PHA or viral antigens, latent and lytic EBV antigens as well as control viral antigens from VZV, Influenza & CMV. Latent and lytic EBV antigen EBNA1, EBNA2, EBNA3A, LMP1, LMP2 and EBV lytic antigen BZLF1 and GP350/450 peptide pools were tested in PBMCs from healthy controls (grey), EBV^-^ DLBCL (black) and EBV^+^ DLBCL (blue), presented as spot-forming cells (SFC) per 10^6^ PBMCs.

### EBV⁺ DLBCL tumours exhibit a relative depletion of cytotoxic T-cells within an expanded TME

While antiviral T-cell immunity was markedly reduced in the blood of DLBCL patients, these findings do not exclude an additional pathogenic contribution of local immunosuppressive mechanisms operating within the tumour. Indeed, systemic and microenvironmental immune dysfunction may converge to facilitate EBV-driven lymphomagenesis. To evaluate this possibility, we performed comprehensive high-dimensional spatial proteomic profiling of EBV^+^ and EBV⁻ DLBCL TMEs using the Phenocycler-FUSION (Akoya Biosciences) platform (Fig. 2a; Supp Table S8). A multiplex immunofluorescence (mIF) panel was used to profile cellular composition and function (Supp. Table S4; Extended Data Fig. 2a,b). Automated cell segmentation identified approximately 1.5 million single cells, which were clustered and annotated according to canonical lineage-defining markers (Fig. 2b,c; Extended Data Fig. 2d). mIF images were cross validated against matched haematoxylin and eosin (H&E) and EBER ISH to confirm spatial integrity and phenotypic fidelity of staining patterns (Fig. 2d). We found that EBV^+^ DLBCL contained significantly more (non-malignant) immune cells and fewer tumour B-cells (TBC) compared with EBV-negative (EBV^-^) tumours (Fig. 2e, f). However, a more detailed phenotypic analysis of the TME revealed that the proportion of CD8^+^ T-cells relative to all immune cells was markedly reduced in EBV^+^ tumours, whereas CD4^+^ T-cell and regulatory T-cell (Treg) frequencies were comparable (Fig. 2g). Together, these data show that despite a quantitatively expanded infiltrate, EBV^+^ DLBCL is relatively deprived of cytotoxic CD8^+^ T-cells. This intratumoural deficit mirrors the marked loss of circulating antiviral T-cell responses we observed in the blood in EBV^+^ DLBCL patients, highlighting a shared deficit in T-cell immunity at both systemic and local levels. Such dual impairment strongly supports a model in which systemic and microenvironmental dysfunction together facilitate EBV-driven lymphomagenesis.

**Figure 2.**
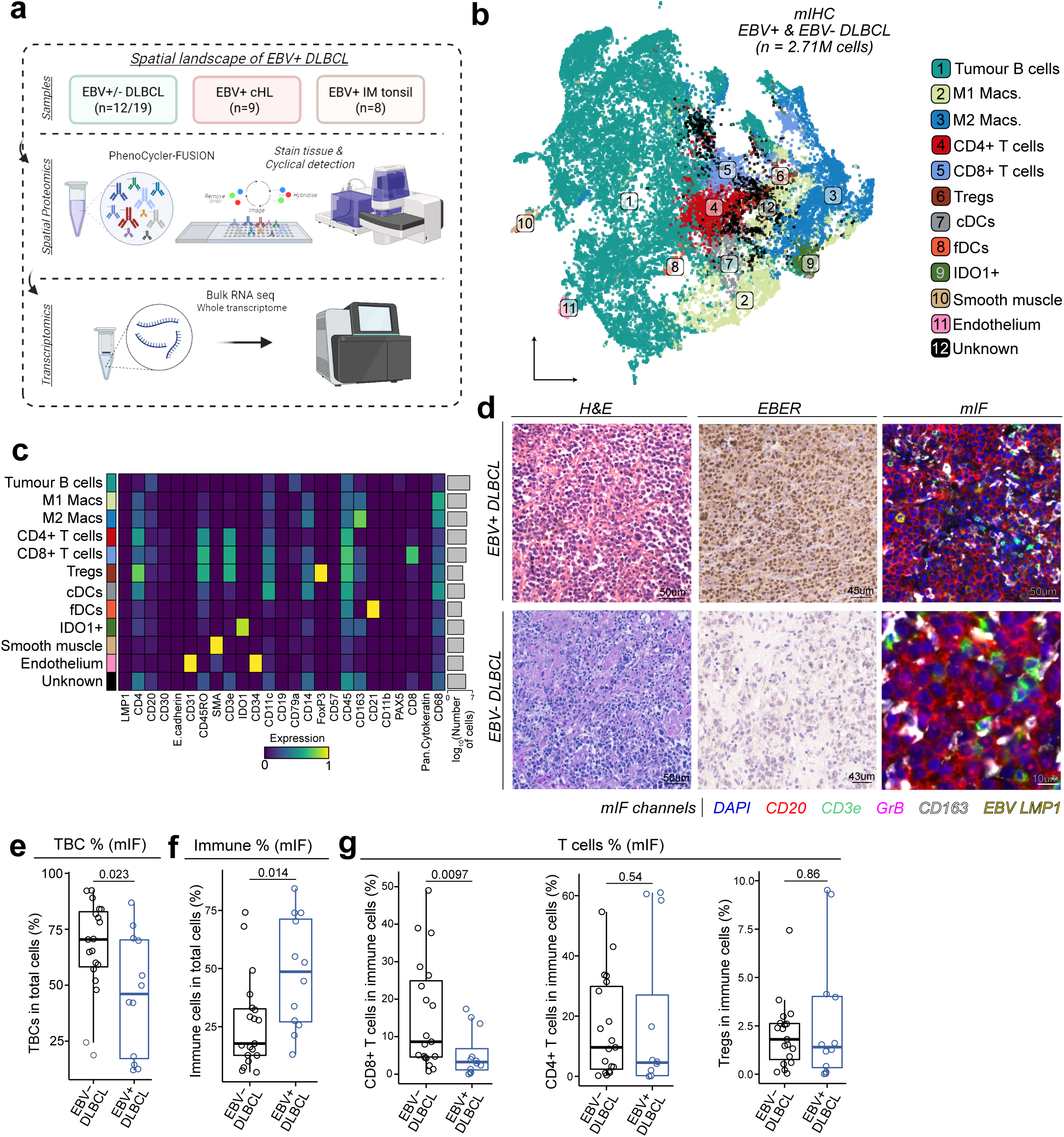
mIF of EBV^+^ DLBCL reveals diminished cytotoxic T-cells. **a.** 12 EBV^+^ DLBCL, 19 EBV^-^ DLBCL, 9 EBV^+^ cHL, and 8 IM tonsils tissues were collected. Tissues were stained and imaged on the Phenocycler-FUSION. RNA was extracted and sequenced. **b.** tSNE of single-cell mIF clustering identifying 12 phenotypes. **c.** Cluster heatmap of 12 phenotypes showing canonical marker expression profiles. **d.** Comparison of H&E, EBER, mIF and cell annotations for representative field of views of EBV^+^ DLBCL and EBV^-^ DLBCL. **e.** Percentage of tumour B-cells in EBV^+^ vs. EBV^-^ DLBCL. **f.** Immune cell abundance in EBV^+^ vs. EBV^-^ DLBCL. **g.** CD8^+^, CD4^+^ and Treg abundances as a function of total immune cells in EBV^+^ vs. EBV^-^ DLBCL.

### A myeloid-dominant immunosuppressive programme is present in EBV⁺ DLBCL tumours

Given that we had observed a relative depletion of intratumoural CD8^+^ cytotoxic T-cells in EBV^+^ DLBCL, we next asked whether a dominant suppressive myeloid programme might be associated with this deficit. We first noted the enrichment of IDO1^+^ M2-like macrophages in EBV^+^ compared to EBV- tumours, despite no increase in total M1 or M2 macrophage abundance (Fig. 3a, b). This skewed balance was functionally reflected in a disproportionate rise in the M2:CD8^+^ T-cell ratio, whereas other macrophage:T-cell ratios remained unchanged (Extended Data Fig. 3a). High-dimensional spatial analysis revealed that the macrophages do not accumulate randomly but organise into distinct immunosuppressive cellular neighbourhoods (see Methods and Extended Data Fig. 3b). Specifically, the IDO1^+^ macrophage/PD-1^+^ Treg-rich neighbourhood (N9) was markedly expanded in EBV^+^ DLBCL, accompanied by a contraction of CD4^+^ and CD8^+^ T-cell-dominated neighbourhoods (N3 and N11) (Fig. 3c, d). Thus, the loss of cytotoxic T-cell niches in EBV^+^ DLBCL is mirrored by the emergence of a structured suppressive myeloid-regulatory T-cell neighbourhood. To investigate the molecular basis of this rewired ecosystem, bulk RNA-seq demonstrated an upregulation of macrophage-associated immunoregulatory genes in EBV^+^ DLBCL, including *SPP1*, *APOO*, *SOX9*, *KLHL21*, and *CCL17*, a chemokine known to drive Treg recruitment (Fig. 3e; Extended Data Fig. 3c)^19^. Conversely, *CCL21*, which is essential for CD4+/CD8+ T-cell trafficking and retention, was downregulated, consistent with impaired CD8+ T-cell access to EBV^+^ tumours^20, 21^. Gene-set enrichment analysis showed that EBV^+^ tumours were enriched for hypoxia-associated pathways which are known to induce SPP1 expression (Fig. 3f). In keeping with a potential role for these immunosuppressive macrophages in driving more aggressive disease, we found that high SPP1 expression was also significantly associated with inferior survival in patients ABC-DLBCL the cell of origin (COO) subtype characteristic of nearly all EBV⁺ tumours (Fig. 3g). Together, these data reveal that EBV^+^ DLBCL harbours a macrophage-dominated immunosuppressive microenvironment in which expansion of IDO1^+^ macrophage niches and SPP1 expression coincides with and likely contributes to the observed reduction in cytotoxic T-cell infiltration.

**Figure 3.**
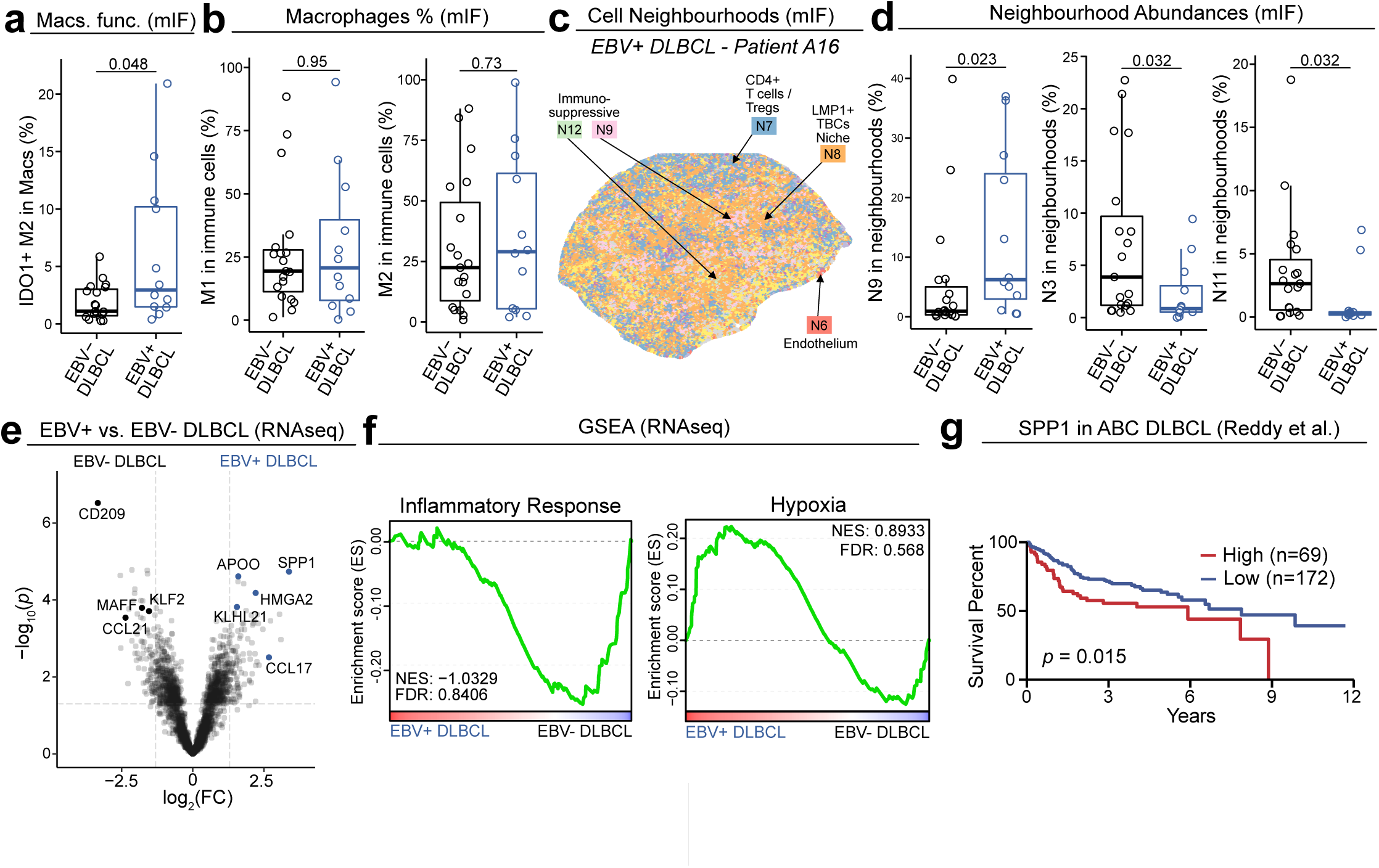
Immunosuppressive macrophages and SPP1 are enriched in the EBV^+^ DLBCL TME. **a.** IDO1^+^ M2 macrophage abundance as a function of total macrophages in EBV^+^ and EBV^-^DLBCL. **b.** M1 and M2 macrophage abundances as a function of total immune cells in EBV^+^ and EBV^-^DLBCL. **c.** Cell neighbourhood visualisation of patient A16 highlighting immunosuppressive, LMP1^+^ tumour, M2 macrophage, cytotoxic and vasculature niches. **d.** Neighbourhood abundances of neighbourhood 9, 3 & 11 in EBV^+^ and EBV^-^ DLBCL. **e.** Differential gene expression volcano plot of EBV^+^ vs. EBV^-^ DLBCL from bulk RNA sequencing data. **f.** Gene set enrichment analysis of EBV^+^ vs. EBV^-^ DLBCL for inflammatory response and hypoxia gene sets. **g.** Survival analysis of SPP1 high vs. low patients in ABC-DLBCL. Value to differentiate high and low patients was calculated by CutOffFinder.

### LMP1 orchestrates an immunosuppressive macrophage niche in EBV^+^ DLBCL

Given the well-established ability of LMP1 to influence both the nature of the EBV-infected cells and the recruitment of immune cells *in vitro*, we next asked whether LMP1 expression in the TBC of primary EBV⁺ DLBCL correlated with shifts in TBC phenotype or with remodelling of the immediate microenvironment. We found that compared with LMP1⁻ TBCs in EBV⁺ DLBCL and with TBCs in EBV^-^ DLBCL, LMP1⁺ TBCs displayed a distinct phenotype, characterised by increased HLA-DR, HLA-A, PD-L1, CD30, CD44 and vimentin expression. CD20 expression was also reduced in LMP1-expressing TBC compared to TBC of EBV- DLBCL (Fig. 4a). To validate that these alterations were LMP1-driven rather than simply correlational, we expressed LMP1 in four EBV^-^ DLBCL cell lines (Extended Data Fig. 4a-c). We found that ectopic LMP1 expression reproduced the same phenotype (Supp. Table S5), including a reduction in CD20 protein expression levels with increasing LMP1 (induced by doxycycline; Fig. 4b,c). We next examined how LMP1 reshapes the broader tumour ecosystem in the primary tumours. We found that LMP1 expression strongly correlated with increased immune-cell infiltration in the TME (Fig. 4d). In EBV⁺ DLBCL, LMP1⁺ TBCs were embedded within dense aggregates of PD-L1⁺ M1-like macrophages, forming highly suppressive tumour-myeloid microdomains (Fig. 4e). PD-L1⁺IDO1⁺ M2-like macrophages were likewise concentrated around LMP1⁺ TBCs but were depleted around LMP1⁻ tumour cells (Extended Data Fig. 4e). These suppressive macrophage configurations were largely absent in EBV-DLBCL (Fig. 4f; Extended Data Fig. 4f,g). Concomitantly, proliferating CD4⁺ and CD8⁺ T-cells were markedly reduced in the immediate vicinity of LMP1⁺ TBC (Fig. 4e). To mechanistically define the signals responsible for these spatially organised suppressive niches, we analysed transcriptional changes induced by LMP1 by performing RNA-seq on the LMP1-expressing DLBCL cell lines as well as EBV-infected EBV^-^ DLBCL cell lines (Extended Data Fig. 4b-d). Both conditions upregulated the monocyte-attractant chemokines *CCL3* and *CCL5* (Fig. 4g), providing a mechanistic explanation for the macrophage-rich microdomains observed *in situ*. Ten genes were consistently upregulated across both conditions, defining a shared LMP1-driven immune-modulatory signature (Extended Data Fig. 4d & Supp. Table S6). Pathway analysis revealed enrichment of activation of inflammatory signalling, chemokine and chemokine activity programmes (Fig. 4h), aligning with the spatial accumulation of PD-L1⁺ and IDO1⁺ macrophages around LMP1⁺ tumour cells. To test whether LMP1 directly influences macrophage phenotype, we exposed THP-1-derived macrophages to supernatants from LMP1-expressing or EBV-infected DLBCL cell lines. Both supernatants induced SPP1 expression in macrophages (Fig. 4i), compared with parental cell line supernatant, revealing that LMP1 can induce SPP1⁺ immunosuppressive macrophages via a paracrine mechanism. Collectively, these data establish LMP1 as a central architect of the EBV⁺ DLBCL microenvironment, reshaping tumour-cell antigen presentation, driving chemokine and immune checkpoint upregulation, recruiting suppressive macrophage populations, and creating IDO1⁺/SPP1⁺ myeloid niches that may inhibit proliferating T-cells and reinforce local immune dysfunction.

**Figure 4.**
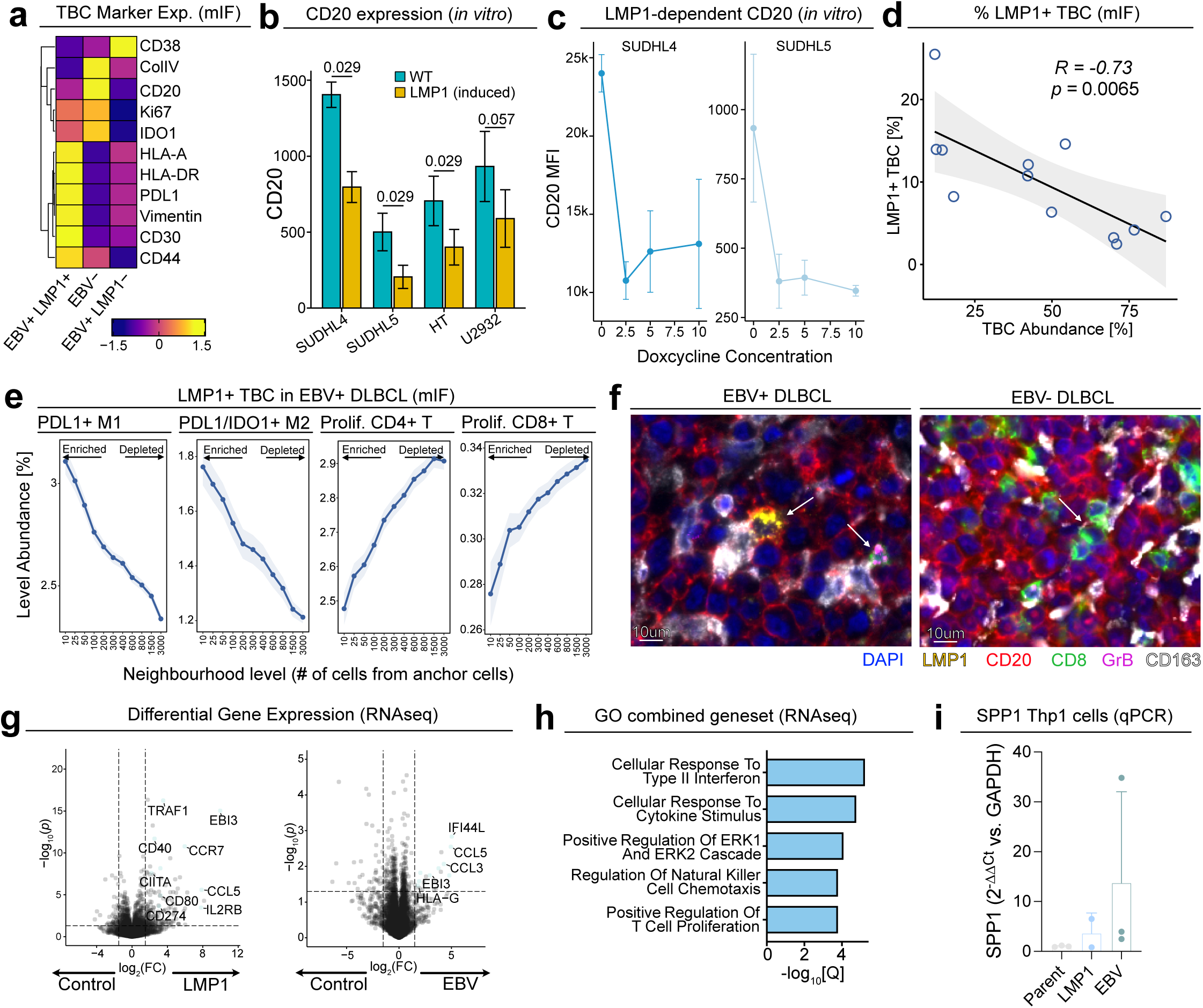
Immediate immune microenvironment of EBV^+^ DLBCL TBCs and macrophages. **a.** Relative expression of functional tumour markers (CD38, Col IV, CD20, Ki67, IDO1, HLA-A, HLA-DR, PDL1, Vimentin, CD30 & CD44) in LMP1^+^/LMP1- EBV^+^ DLBCL TBCs and EBV- DLBCL TBCs. **b.** CD20 expression in parent and LMP1 induced SUDHL4, SUDHL5, U2932 and HT DLBCL cell lines. **c.** CD20 expression as a function of doxycycline inducible LMP1 expression in SUDHL4 and SUDHL5 DLBCL cell lines. **d.** Percent of LMP1^+^ tumour B-cells (of all tumour B-cells) in each patient as a function of total tumour B-cells. **e.** TME quantification (cell abundance as a function of distance) of LMP1^+^ TBCs for PDL1^+^ M1 macrophages, PDL1^+^IDO1^+^ M2 macrophages, proliferative CD4^+^ T-cells and proliferative CD8^+^ T-cells. **f.** Representative mIF images of different TMEs of EBV^+^ and EBV^-^ DLBCL. **g.** Differential gene expression of LMP1 transfected cells vs. control and EBV infected cells vs. control. Differential expression testing was conducted with the DLBCL COO classification as a co-variate in DESeq2^37^. **h.** GO biological processes pathway analysis of commonly upregulated genes. **i.** qPCR quantification of SPP1 in differentiated THP-1 cells after culture in DLBCL cell line supernatant.

### EBV^+^ DLBCL exhibits a uniquely suppressive immune architecture compared with EBV-associated classical Hodgkin lymphoma and infectious mononucleosis

Given that EBV is associated with B-cell lymphoproliferative diseases, we next examined whether the profoundly immunosuppressive features of the EBV⁺ DLBCL TME we observed were disease-specific or represented a more general phenomenon. To do this, we directly compared the spatial proteomic profiles of EBV⁺ DLBCL with two other EBV-driven B-cell lymphoproliferations; EBV⁺ classical Hodgkin lymphoma (cHL) and symptomatic primary benign EBV infection (infectious mononucleosis; IM) using the same mIF panel (Extended Data Fig. 5a)^22^. Across all cohorts, EBV⁺ DLBCL was consistently the most lymphocyte-depleted state, exhibiting markedly reduced CD8⁺ and CD4⁺ T-cell abundances relative to both cHL and IM (Fig. 5a; Extended Data Fig. 5b). In contrast, compared with cHL, M1 macrophages were significantly increased in EBV^+^ DLBCL. Moreover, while overall M2 macrophage numbers were similar across disease states (Fig. 5b), compared with cHL, M2 macrophages in EBV⁺ DLBCL expressed substantially higher levels of HLA-DR, indicating enhanced activation and antigen-presenting capacity (Fig. 5c). Notably, the T-cells in EBV⁺ DLBCL also displayed a uniformly activated/exhausted phenotype, characterised by markedly elevated PD-1 expression across CD4⁺, CD8⁺ and Treg populations (Fig. 5c; Extended Data Fig. 5c), together with reduced proliferative capacity compared with IM (Extended Data Fig. 5c). This constellation of lymphocyte depletion, PD-1 upregulation and macrophage enrichment closely mirrors the differences we observed between EBV⁺ and EBV⁻ DLBCL, positioning EBV⁺ DLBCL at the extreme end of an immunosuppressive continuum. To understand the basis of these cross-disease differences, we next examined the organisation of LMP1-centred immune niches, reasoning that this EBV-encoded oncoprotein might impart disease-specific immunological programmes. Spatial profiling revealed striking variation in LMP1-associated macrophage configurations across the three diseases (Fig. 5d; Extended Data Fig. 5a,c,d). In EBV⁺ DLBCL, PD-L1⁺ M1-like macrophages densely surround LMP1⁺ TBCs, forming suppressive microdomains that were also enriched for PD-1⁺ T-regs and exhausted T-cells. In contrast, in EBV⁺ cHL, PD-L1⁺ M2-like macrophages predominate around LMP1⁺ Hodgkin/Reed-Sternberg cells. In IM, PD-L1⁺IDO1⁺ M1-like macrophages aggregate around EBV-infected B-cells, reflecting an acute antiviral response rather than the chronic suppressive configuration observed in malignancy. These comparisons demonstrate that although LMP1 expression is shared across these EBV-associated diseases, its impact on the composition appear context-dependent. Importantly, its most suppressive effects occur in EBV⁺ DLBCL, where LMP1 expression is associated with dense PD-L1⁺ macrophage aggregates, enrichment of PD-1⁺ Tregs and exhausted T-cells (Fig. 5e). Taken together, these data identify EBV⁺ DLBCL as the most immunosuppressed and macrophage-dominated entity among these EBV-associated lymphoproliferative disorders.

**Figure 5.**
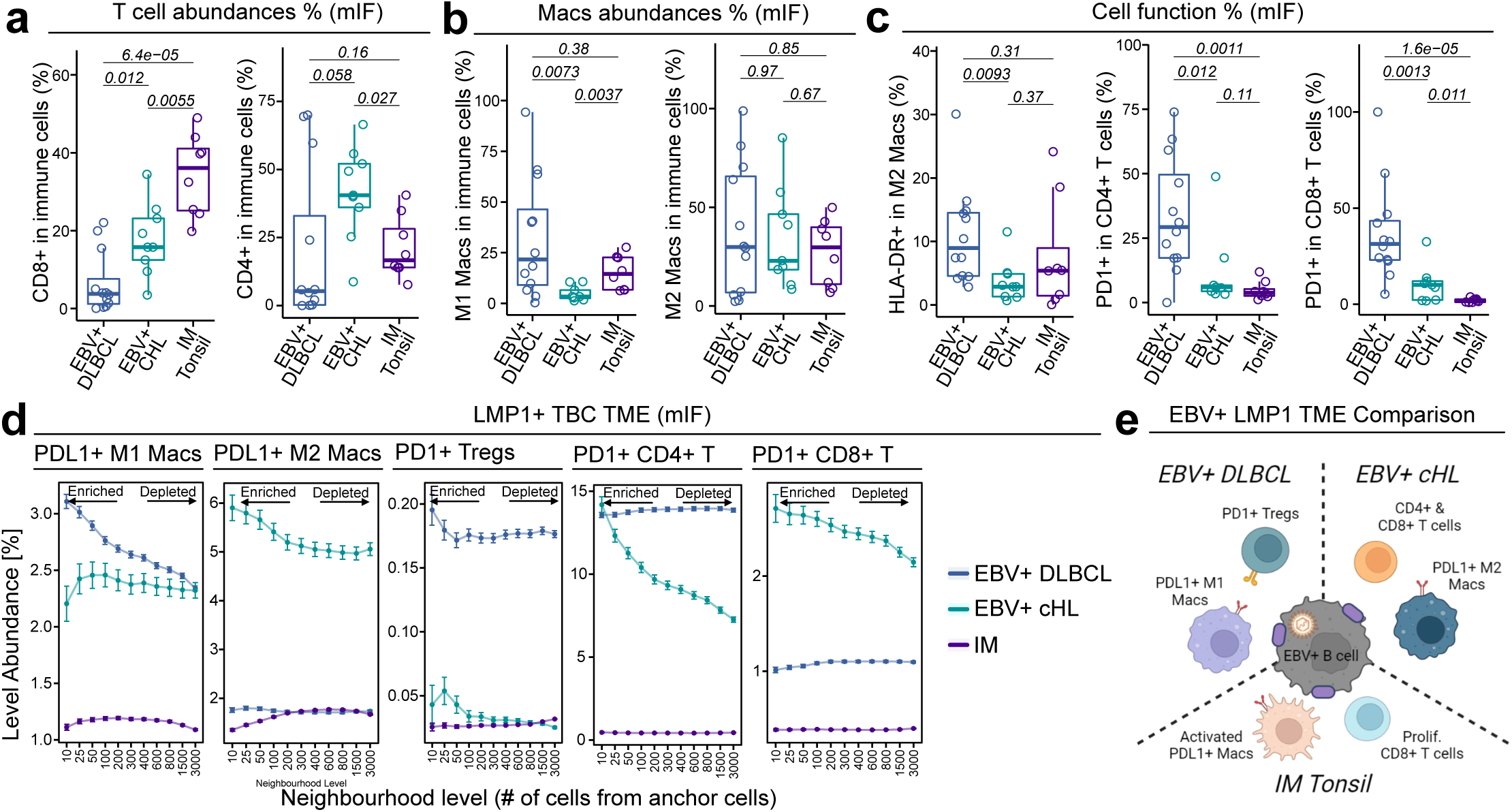
The EBV^+^ DLBCL TME is more suppressive than EBV^+^ cHL and IM Tonsil. **a-b.** CD8^+^ T-cells and CD4^+^ T-cells (a) and M1 macrophages and M2 macrophages (b) abundances as a function of total immune cells in EBV^+^ DLBCL, EBV^+^ cHL and IM Tonsil. **c.** HLA-DR^+^ M2 Macrophages, PD1^+^ CD4^+^ T-cells and PD1^+^ CD8^+^ T-cells abundances as a function of total immune cells in EBV^+^ DLBCL, EBV^+^ cHL and IM Tonsil. **d.** TME quantification (cell abundance as a function of distance) of LMP1^+^ TBCs/HRS cells/EBV-infected B-cells for PDL1^+^ M1/M2 macrophages, PD1^+^ Tregs and exhausted CD4^+^/CD8^+^ T-cells. Kruskal-Wallis test was used at each TME level. **e.** Graphic illustration of the TME differences between EBV^+^ DLBCL, EBV^+^ cHL & IM Tonsil.

## Discussion

EBV^+^ DLBCL has long been viewed as a disorder arising from immunosenescence, with impaired viral immunity in older adults proposed as a permissive state for EBV-driven lymphomagenesis^2, 3, 23^. However, the recognition that EBV⁺ DLBCL arises across all age groups, including younger and ostensibly immunocompetent individuals, has called into question this view^7, 8^. By integrating functional profiling of systemic T-cell immunity, high-plex spatial proteomics, and mechanistic *in vitro* modelling, we reveal that EBV⁺ DLBCL is underpinned not by a single dominant defect but by the convergence of reduced systemic antiviral T-cell responses and a highly orchestrated, LMP1-driven immunosuppressive TME.

We found that patients with DLBCL exhibit broad impairments in circulating memory T-cell responses across a wide range of viral antigens, including EBV latent and lytic proteins, with the most profound deficits observed in the EBV⁺ DLBCL group. We also showed that T-cell responses to antigens from unrelated viruses such as influenza and VZV were similarly reduced, indicating that this dysfunction reflects a generalised collapse of antiviral memory T-cell immunity rather than an EBV-restricted defect. Importantly, the magnitude of these impairments was comparable in both younger and older patients, underscoring that systemic immune dysfunction in DLBCL cannot be attributed solely to immunosenescence or age-related decline. Whether this systemic dysfunction precedes lymphoma development or is secondary to the metabolic and inflammatory burden of malignancy remains unresolved. Nevertheless, reduced antiviral surveillance may contribute to inadequate control of EBV lytic reactivation, consistent with prior reports linking EBV DNA load to poorer outcomes in EBV^+^ lymphomas, including EBV^+^ DLBCL^6, 24–26^.

These broad reductions in antiviral T-cell responses in all the EBV^+^ DLBCL patients we tested prompted us to examine whether a complementary form of immune dysfunction might also be operating within the TME. Despite evidence of profoundly reduced systemic antiviral T-cell responses, EBV⁺ DLBCL tumours overall contained more immune cells than EBV⁻ cases. However, high-dimensional spatial analysis revealed that these infiltrates are qualitatively abnormal since the EBV⁺ tumours are markedly depleted of cytotoxic CD8⁺ T-cells and contain expanded populations of PD-1⁺ CD4⁺ T-cells and regulatory T-cells compared with EBV-tumours. This dual deficit, systemic and local, points to a failure both to generate effective antiviral T-cell immunity in the blood, and to recruit or maintain cytotoxic T-cells within the tumour.

One of the most striking findings was the observation that EBV⁺ DLBCL is characterised by a prominent myeloid-dominated immunosuppressive programme in which IDO1⁺ M2-like macrophages are significantly enriched, forming spatially coherenT-cellular neighbourhoods with PD-1⁺ Tregs and exhausted T-cells. These suppressive niches were associated with the contraction of cytotoxic neighbourhoods and may serve as key gatekeepers that restrict effector T-cell access. Mechanistically, IDO1 activity drives tryptophan catabolism and kynurenine accumulation and impaired T-cell proliferation and function^27–29^, providing a plausible explanation for how these macrophage-rich niches enforce local immune suppression. Moreover, bulk RNA-sequencing revealed that EBV⁺ tumours upregulate a macrophage-immunoregulatory signature, that includes *SPP1*, *CCL17*, *KLHL21*, *SOX9*, as well as other genes associated with immunosuppressive macrophage differentiation^30, 31^. *SPP1* expression has recently been identified as a prognostically adverse factor across multiple cancers. We found that high SPP1 expression was associated with inferior overall survival in ABC-DLBCL, the COO subtype most commonly represented in EBV⁺ DLBCL^32, 33^. Mechanistically, supernatants from LMP1-expressing or EBV-infected DLBCL cell lines directly induced SPP1 expression in THP-1 macrophages, demonstrating a tumour-derived paracrine link between EBV infection and the emergence of SPP1⁺ macrophages. Together with our validation of marked SPP1⁺ macrophage enrichment in an independent large FFPE cohort of EBV^+^ DLBCL, these findings identify SPP1 as a central and clinically relevant feature of the EBV⁺ DLBCL microenvironment. Although SPP1 is known to be upregulated by platelet-derived factors and hypoxic niches^32, 33^, our results raise the possibility that hypoxia-driven macrophage programmes may also contribute to DLBCL pathogenesis, potentially acting in parallel with EBV-mediated induction of SPP1.

LMP1 expression, present in all our EBV⁺ cases although in variable frequencies of TBCs for each patient, emerged as a major organiser of the tumour ecosystem. LMP1⁺ tumour B-cells displayed hallmarks of immune modulation: upregulation of HLA-A, HLA-DR, PD-L1, CD30, CD44, and vimentin, and downregulation of CD20 – as shown previously^34^ – validated through inducible LMP1 expression in multiple DLBCL cell lines. This latter phenotype may directly contribute to inferior rituximab responses observed in EBV⁺ disease^26^, but this requires further confirmation. Spatially, LMP1⁺ tumour clusters were surrounded by dense cuffs of PD-L1⁺ and IDO1⁺ macrophages, with simultaneous exclusion of proliferating CD4⁺ and CD8⁺ T-cells and enrichment of PD-1⁺ Tregs. RNA-sequencing of LMP1-transduced and EBV-infected DLBCL cell lines identified a shared signature, which included the upregulation of the potent monocyte-attractant chemokines *CCL3* and *CCL5*^35, 36^, providing a mechanistic explanation for the macrophage-rich microdomains observed in the tumour samples. Collectively, these findings define LMP1 as a central driver of the immunosuppressive TME of EBV^+^ DLBCL.

Comparisons with EBV⁺ cHL and IM revealed that EBV⁺ DLBCL exhibits the most extreme degree of immune depletion and suppressive myeloid organisation among these EBV-associated lymphoproliferative disorders. EBV⁺ DLBCL also contained the lowest relative frequencies of intratumoural T-cells, the highest positivity of PD-1 expression on both CD4⁺ and CD8⁺ populations, and the most pronounced enrichment of PD-L1⁺ macrophages. Notably, LMP1-associated microenvironments also differed markedly across diseases. In EBV⁺ DLBCL, PD-L1⁺ M1-like macrophages and IDO1⁺ M2-like macrophages formed dense suppressive rings around tumour cells, whereas EBV⁺ cHL showed a predominance of PD-L1⁺ M2-like macrophages but with more preserved lymphocyte infiltration. By contrast, IM exhibited macrophage aggregates typical of acute antiviral inflammation rather than chronic immunosuppression. These disease-specific patterns highlight that the immunological imprint of LMP1 is profoundly context-dependent.

In conclusion, this study provides the most comprehensive immunological characterisation of EBV⁺ DLBCL to date, demonstrating that its pathogenesis arises from the convergence of systemic antiviral immune failure and a highly structured, LMP1-driven, myeloid-dominated TME. These findings not only redefine the immunobiology of EBV⁺ DLBCL, but they also highlight the tumour-macrophage interface as a promising therapeutic target in this poor-prognosis lymphoma. In particular, strategies that disrupt the myeloid-tumour axis, such as the inhibition of IDO1, targeting of SPP1, blockade of CCL3/CCL5-mediated monocyte recruitment, or myeloid-reprogramming agents such as CSF1R inhibitors, may help overcome the dominant immunosuppressive niches characteristic of this disease.

## Extended Data Figure Legends

**Extended Data Figure 1.**
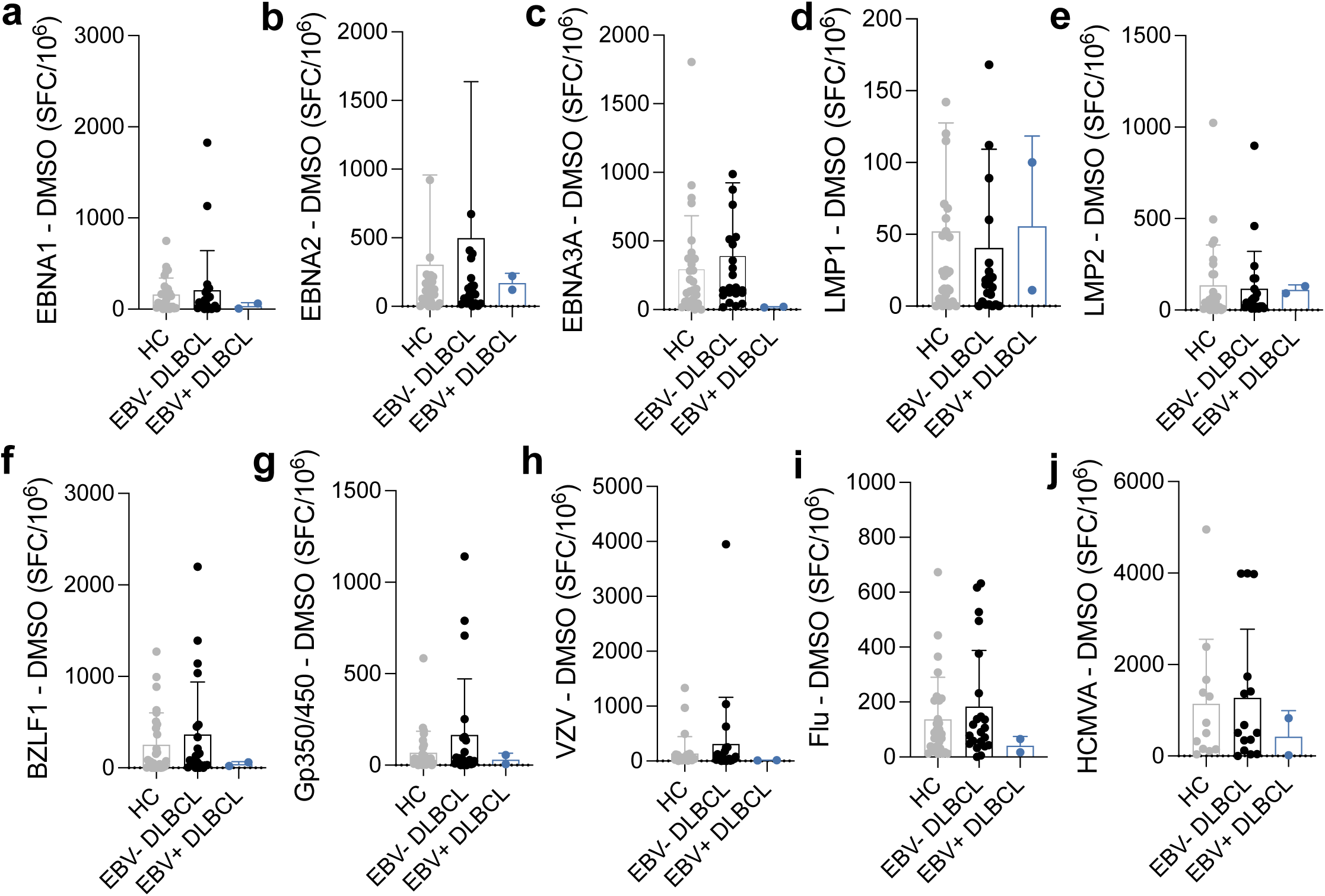
Frequency of T-cells specific for viral antigens in EBV^+^ and EBV^-^DLBCL patients compared with healthy controls (validation cohort) **a - j.** IFNγ ELISpot response to latent and lytic EBV antigens as well as control viruses (VZV, Influenza & CMV). Latent and lytic EBV antigen EBNA1, EBNA2, EBNA3A, LMP1, LMP2 and EBV lytic antigen BZLF1 and GP350/450 peptide pools were tested in PBMCs from heathy controls (grey), EBV^-^ DLBCL (black) and EBV^+^ DLBCL (blue), presented as spot-forming cells (SFC) per 10^6^ PBMCs.

**Extended Data Figure 2.**
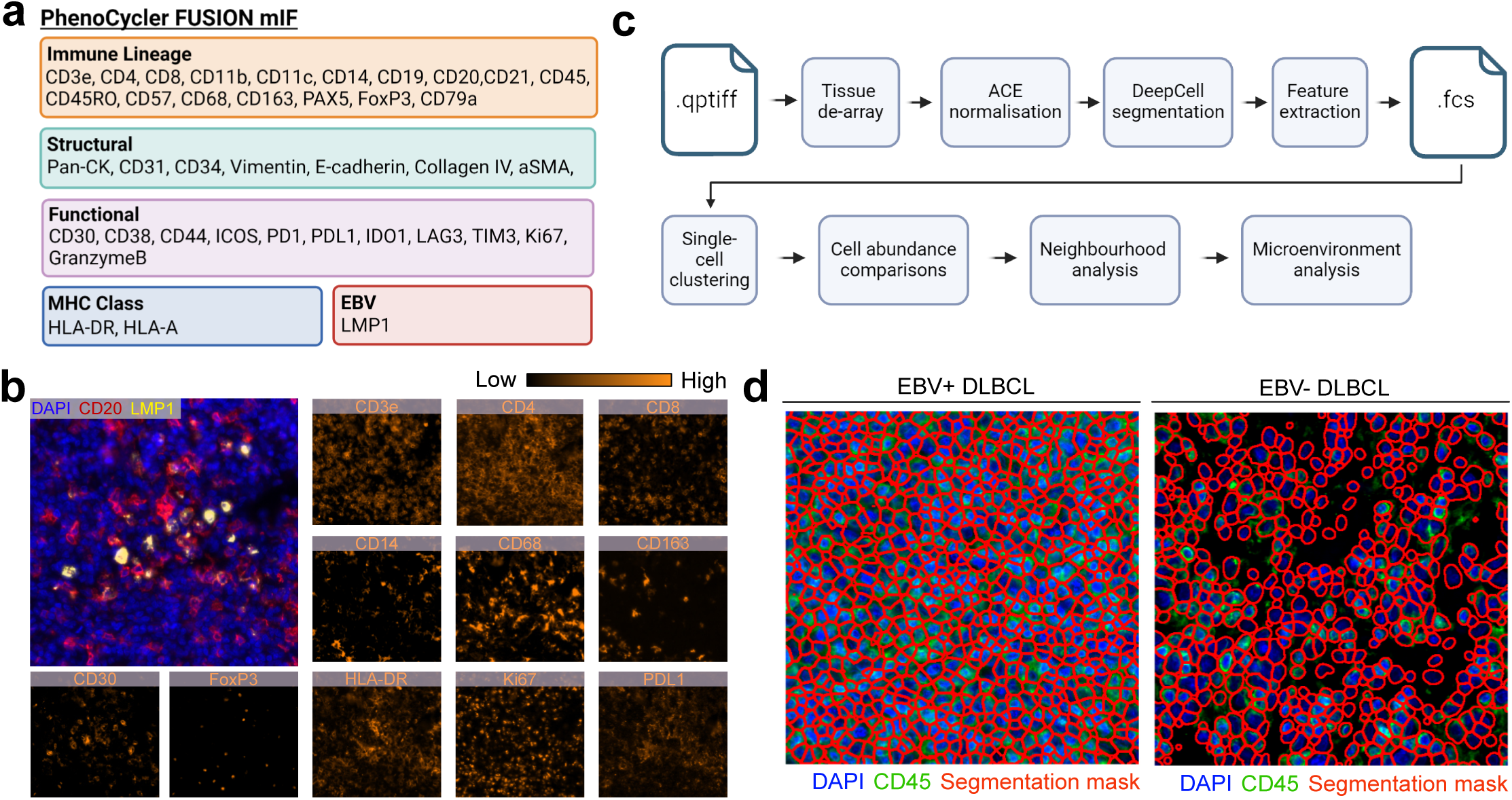
Overview of mIF panel, image pre-processing, and segmentation. **a.** Phenocycler-FUSION antibody panel **b.** Example Phenocycler-FUSION staining of CD3e, CD4, CD8, CD14, CD68, CD163, CD30, FoxP3, HLA-DR, Ki67 and PDL1 from a representative field of view of EBV^+^ DLBCL. **c.** mIF image pre-processing and analysis pipeline used for Phenocycler-FUSION images. **d.** Examples of single-cell segmentation using DeepCell.

**Extended Data Figure 3.**
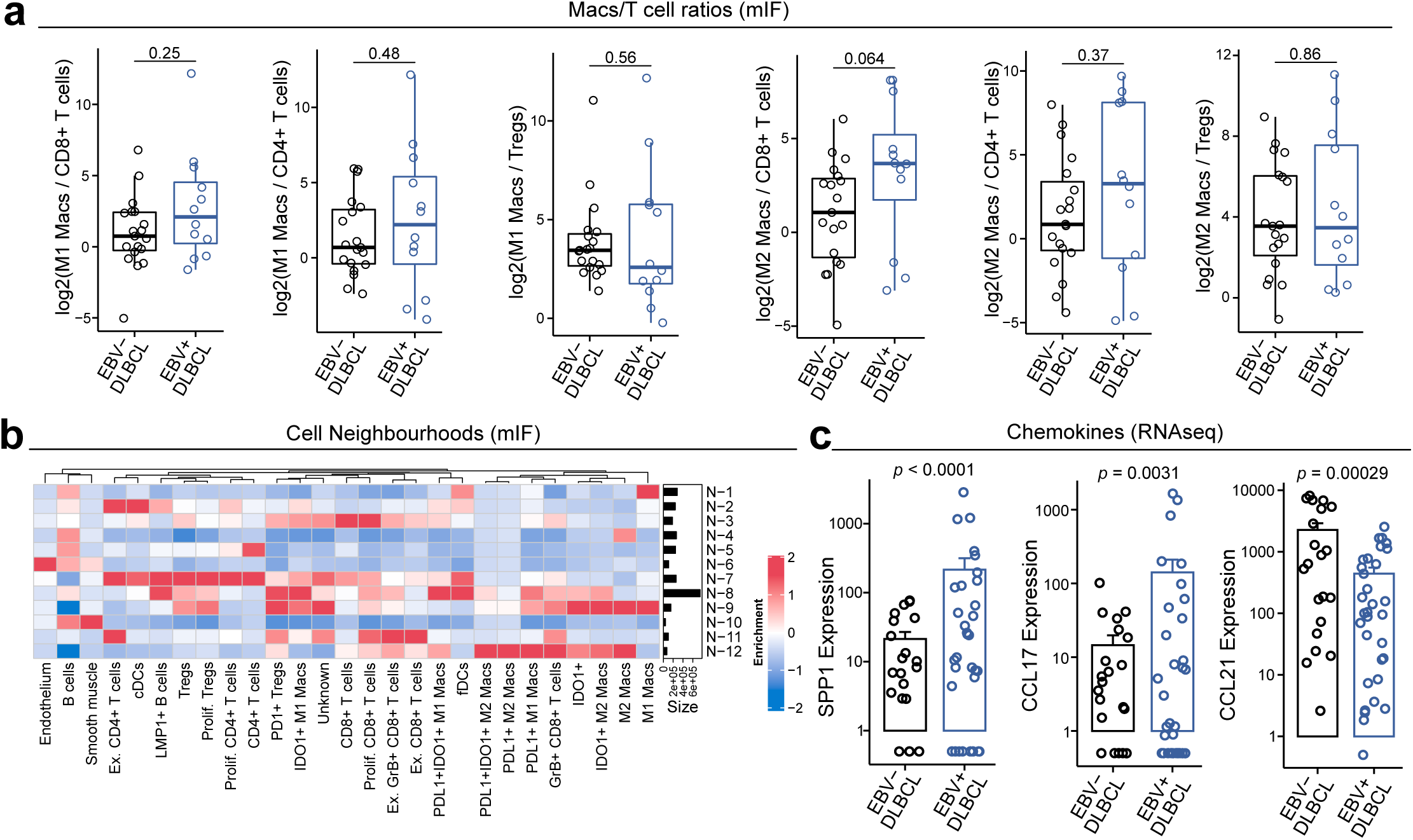
Additional EBV^+^ TME data. **a.** Ratios of Macrophage and T-cell phenotypes between EBV^+^ and EBV^-^ DLBCL. **b.** Cellular neighbourhood cell enrichment heatmap showing the relative abundances of cell types within each neighbourhood. **c.** Chemokine levels from bulk RNAseq analysis in EBV^+^ vs. EBV^-^ DLBCL.

**Extended Data Figure 4.**
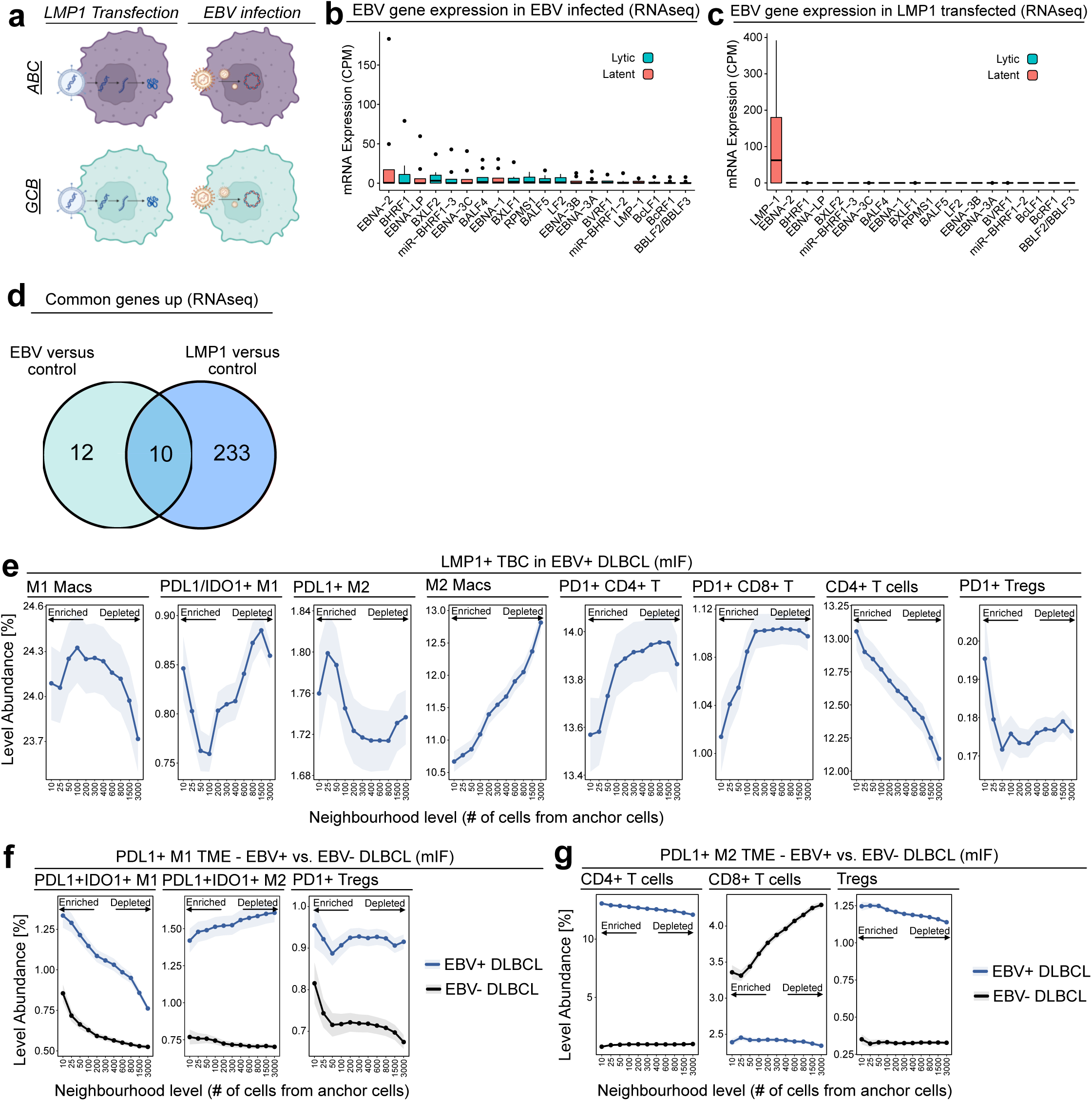
Spatial niches of macrophages and LMP1^+^ TBCs in EBV^+^ DLBCL. **a.** LMP1 transfection and EBV infection of four human DLBCL cell lines (SUDHL4, SUDHL5, HT & U2392). **b.-c.** EBV gene expression of (e) EBV infected and (f) LMP1 transfected hDLBCL cell lines. **d.** Commonly upregulated genes in LMP1 transfected and EBV infected DLBCL cell lines. **e.** TME quantification (cell abundance as a function of distance) of LMP1^+^ TBCs for M1 macrophages, PDL1^+^IDO1^+^ M1 macrophages, PDL1^+^ M2 macrophages, M2 macrophages, exhausted CD4^+^ T-cells, exhausted CD8^+^ T-cells, CD4^+^ T-cells and PD1^+^ T regs. **f. & g.** TME quantification (cell abundance as a function of distance) of (b) PDL1^+^ M1 macrophage for PDL1^+^IDO1^+^ M1 macrophages, PDL1^+^IDO1^+^ M2 macrophages and PD1^+^ Tregs and (c) PDL1^+^ M2 macrophages for CD4^+^ T-cells, CD8^+^ T-cells and Tregs. Kruskal-wallis test was used at each TME level.

**Extended Data Figure 5.**
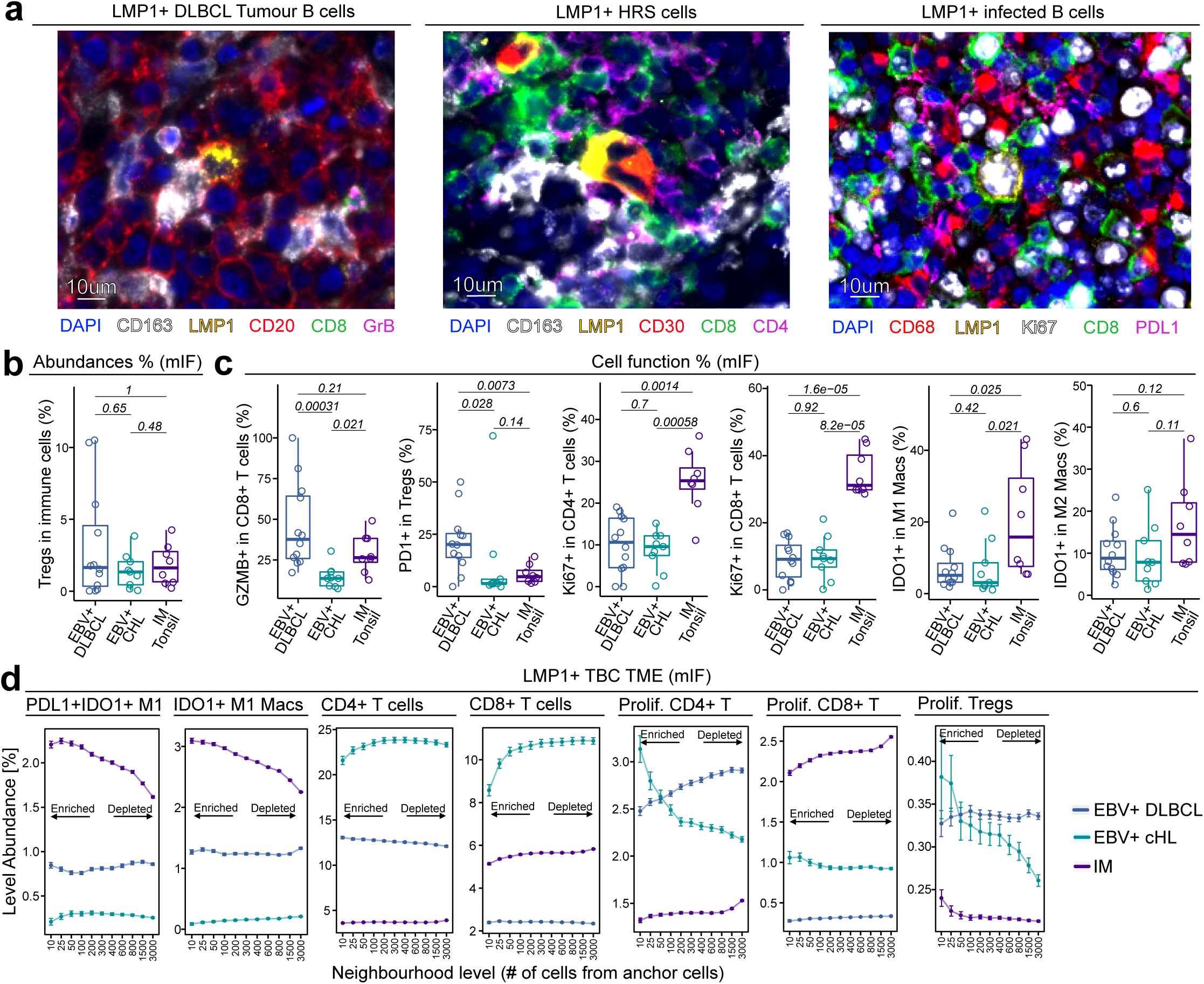
EBV^+^ disease- mIF images and microenvironment. **a.** Representative mIF images of different TMEs of EBV^+^ DLBCL TBCs, EBV^+^ cHL HRS cells and EBV-infected B-cells in IM Tonsil. **b.** Treg abundances as a function of total immune cells in EBV^+^ DLBCL, EBV^+^ cHL & IM Tonsil. **c.** GranzymeB^+^ CD8^+^ T-cell, PD1^+^ Treg, Ki67^+^ CD4^+^ T-cell, Ki67^+^ CD8^+^ T-cell, IDO1^+^ M1 macrophage and IDO1^+^ M2 macrophage abundances as a function of total immune in EBV^+^ DLBCL, EBV^+^ cHL & IM Tonsil. **d.** TME quantification (cell abundance as a function of distance) of LMP1^+^ TBCs for PDL1^+^IDO1^+^ M1 macrophages, IDO1^+^ M1 macrophages, CD4^+^ T-cells, CD8^+^ T-cells, proliferative CD4^+^ cells, proliferative CD8^+^ T-cells and proliferative Tregs.

## Methods

### Blood samples ethical statements, collection & preparation

The EBV biomarkers study was approved by the Metro South Hospital and Health Service (Brisbane, Queensland, Australia; HREC/07/QPAH/035), and all donors provided written informed consent. The ICEPOP study (validation cohort) was approved by the West Midlands (Black Country) Research Ethics Committee (study reference 15/WM/0419) all donors provided written informed consent. Blood samples from DLBCL patients and healthy age and sex matched volunteers from the community were collected in EDTA (for plasma) and Lithium Heparin (LiHep, for PBMC) tubes and prepared for long term storage in liquid nitrogen.

### EBV and CMV serostatus

CMV and EBV serostatus was determined in all donors using an Anti-CMV-IgG and an Anti-EBV-Capsid Antigen-IgG ELISA kit (Euroimmune). Assays were conducted following the manufacturer’s instructions and results determined in respect of the included control reagents.

### EBER ISH

In some cases, access to the diagnostic biopsy FFPE blocks of ICEPOP DLBCL patients was used to corroborate EBV^+^ DLBCL diagnoses. 5um sections were cut from suspected EBV^+^ DLBCL patient FFPE blocks. Slides were dewaxed in Histoclear II, and rehydrated. Target unmasking by boiling in citrate buffer (4.3mM sodium citrate, 1.3mM citric acid, pH 6), 20mins. Slides were washed in RNase free water and dipped in 96% EtOH and briefly air dried. FITC-conjugated EBER probe (Dako) was applied and incubated at 55°C for 90mins. Slides were then washed in stringent wash solution (Dako, 15min 55°C) and TBS-T (Tris buffered saline ^+^ 0.1% Tween 20, Thermo scientific). Anti-FITC-alkaline phosphatase conjugated secondary (Dako) was then applied, 35min RT. Following washing with TBS-T, signal was developed with BCIP (Dako), and counterstained with Vector Red (Vector Labs), dehydrated and mounted with DPX.

### IFNγ ELISpot T-cell responses

Cryopreserved PBMCs were thawed in a water bath at 37°C, diluted with 10ml R10 (RPMI supplemented with 10% batch tested fatal calf serum, and penicillin/streptomycin) and spun at 350g for 10 mins. Cell pellets were resuspended in R10 and cells rested overnight in a humidified incubator (37°C 5% CO2) prior to testing in IFNγ ELISpot assays (Mabtech). Overlapping 15mer peptide pools (detailed in Supp. Table S3) were purchased from JPT Peptide Technologies. All peptides were dissolved in dimethyl sulfoxide (DMSO, Sigma), aliquoted and stored at −20°C. Peptide was added to wells at a final concentration of 1μg/ml. DMSO alone, and Phytohemagglutinin (10μg/ml) were included as negative and positive controls respectively. IFNγ ELISpot assays were as previously described^38^. Briefly 96-well polyvinylidene difluoride-backed plates (Millipore, Bedford, Mass.) were precoated with anti-IFNγ antibody (clone 1-DIK, (MABTECH, Stockholm, Sweden) 15μg/ml) overnight at 4°C. The next day PBMCs were counted using a disposable hemacytometer with a minimum of two separate cell suspension aliquots counted. Typically, 3×10^5^ cells were added per well with each antigen tested in triplicate. Cells were cultured in the ELISpot plate overnight at 37°C in 5% CO2. Secreted IFNγ was detected with a biotinylated anti-IFNγ antibody, (clone 7-B6-1 (Mabtech), 1 μg/ml), followed by streptavidin-conjugated alkaline phosphatase (Mabtech).

Spots were visualised using an alkaline phosphatase substrate kit (Bio-Rad) and counted using an automated plate counter (AID) with AID ELISpot software version 6. Final counts were normalised to the DMSO negative control wells, by subtraction. In all experiments, results from ELISpot assays are expressed as spot-forming cells (SFC) per million PBMCs.

### Tissue cohorts & preparation

Primary DLBCL tumours were collected in Cardiff (Wales; n = 14) and Siena (Italy; n = 11) under their respective ethics. The tissue was formalin-fixed and paraffin-embedded (FFPE). Cases were diagnosed based on Haematoxylin & Eosin (H&E) and Giemsa-stained sections as well as immunophenotyping to determine the cell of origin according to the Hans algorithm. EBV^+^ DLBCL was diagnosed if >80% of cells tested positive by EBER staining, in accordance with the 4th edition of the WHO classification and the International Consensus Classification (ICC). For Phenocycler-FUSION mIF staining, 3mm tissue sections were extracted from blocks and embedded into a tissue microarrays in an arrangement which covered a 27 x 15mm window (in some cases two TMAs were cut onto a single slide). Slides were cut onto positively charged frost-free slides in preparation for staining. Curls were also cut in preparation for RNA extraction and sequencing.

### Chromogenic IHC

FFPE slides were baked for 2 hours prior to staining to ensure yssue adherence. The Leica Bond RXm IHC protocol (DAB) was used with the following condiyons: bake and de-wax, epitope retrieval with a pH 9 buffer for 20 minutes and primary anybody incubayon for 1 hour.

Slides were subsequently mounted with DPX mounyng media and scanned on the FUSION microscope.

### Phenocycler FUSION multiplex immunohistochemistry staining & imaging

A 38-plex Phenocycler FUSION panel was designed, validated and deployed at Akoya Biosciences, Menlo Park, California, USA. The panels, clones and barcodes used are available in Supp. Table S4. Glass slides were prepared, stained, and fixed as per the PhenoCycler protocol. Briefly, slides were deparaffinised in Xylene and re-hydrated in decreasing concentrations of ethanol (100%, 90%, 70% & 50%). Heat Induced Epitope Retrieval (HIER) was performed in a pressure cooker at 110°C for 18 minutes in Tris-EDTA buffer (pH = 9.0). The tissue was then incubated in PhenoCycler Staining Buffer (Akoya Biosciences) for 30 minutes to block non-specific binding of antibodies. Subsequently, the tissue was incubated in a cocktail of the conjugated antibodies overnight at 4°C, then fixed in 4% paraformaldehyde for 10 minutes, 100% methanol for 5 minutes, Fixative Reagent (Akoya Biosciences) for 20 minutes and stored until imaging. Akoya Reporters were added to the corresponding well of a 96-well plate in preparation for imaging, based on the cycle design of the experiment. Autofluorescence subtraction, stitching and compression were completed in the FUSION software resulting in .qptiff files in preparation for processing and analysis.

### PhenoCycler FUSION pre-processing & segmentation

TIFF images were processed through CellMAPS by de-arraying, normalising, pseudo-membrane marker generation, cellular segmentation, and feature extraction^39^. CD45, CD31 and Pan-cytokeratin channels were merged to create a pseudo-membrane marker for segmentation. Nuclear- and membrane-based segmentation was completed on the DAPI and pseudo-membrane marker channels using DeepCell^40^. Segmentation masks and their borders for each core were saved. FCS files were generated using the segmentation masks with the ‘regionprops‘ function in Python.

### Cell type annotation

FCS files generated from Phenocycler-FUSION imaging were imported into MISSILe. Cells with a total size in the top and bottom 0.01% quantile were removed from the analysis. Clustering was performed with FlowSOM with lineage markers used as input. Heatmaps for the enrichment and expression of each marker per cluster were constructed and clusters were merged where necessary and annotated. Tumour B-cells were denoted as any B-cell marker (e.g. CD20, PAX5, CD79B) positive cell. Clusters were qualitatively validated by plotting every cell coloured by phenotype and comparing them to the original staining patterns. For functional markers, cells were assigned positive based on the results of the findCutoff function for markers in each region^41^. Abundances of each cluster were calculated as a percentage of total detected cells per region.

### Tissue neighbourhoods

Cellular neighbourhoods were generated, as previously described^42^, by calculating the 10 nearest neighbours (RANN library) of each cell and clustering these using k-Nearest Neighbours (kNN) with k=12. Neighbourhoods were annotated according to their cell composition.

### Microenvironment analysis

To quantify the immediate cellular microenvironment of LMP1^+^ TBCs (anchor cell), cell proportions were calculated with increasing distances from anchor cells. The nn2 function (RANN library) calculated the N (where N = 10, 20, 40, 80, 100, 200, 400, & 800) number of cells closest to anchor cells. The proportion of each phenotype was then calculated as a percentage of the total cells in each bin (N) and plotted.

### DLBCL cell lines

The DLBCL cell lines SUDHL4 (GCB), SUDHL5 (GCB), HT (GCB) and U2932 (ABC) were maintained in RPMI^+^10% FBS^+^1%P/S/G (Sigma). The U2932 cell lines were flow sorted for CD21 expression. All cell lines were infected/transduced with viruses kindly provided by Dr. Claire Shannon-Lowe, University of Birmingham.

### EBV infection of DLBCL cell lines

Cell lines were first assessed for expression of CD21 (required for EBV infection) by flow cytometry, using anti-CD21-PE-Cy5 (BD Bioscience). Lines without CD21 (SUDHL-4 and -5) were transiently transduced with a CD21 lentivirus construct, CD21 expression was verified by flow cytometry 24-72hrs after transduction. Lines were then exposed to purified Akata strain EBV virus, this virus contains a GFP transgene insert and Neomycin resistance gene for selection, the former allowed identification of infected cells. Briefly 2×10^5^ cells were pelleted and resuspended in concentrated Akata virus in a final volume of 250-500ul. Cells were left in a 37°C incubator to infect overnight. Cells were then cultured and infection assessed by flow cytometry. GFP^+^ cells were FACS sorted, however long-term stability of EBV infection was found to be poor, as such cells were standardly maintained in empirically determined G418 concentrations resulting in stable EBV infection.

### LMP1 transduction of DLBCL cell lines

Cell lines were transduced with lentivirus containing the LMP1 gene under a Dox-on promoter and GFP transgene for selection. Cells were exposed to lentivirus in the presence of 5ug/ml polybrene for 3-5hrs at 37°C. Cells were then cultured and transduction determined by flow cytometry. To produce a pure LMP1 transduced population cells were FACS sorted, cultured and then resorted resulting in pure LMP1 transduced populations. Cells were maintained in the absence of doxycycline.

### RNA extraction & sequencing of cell lines

Prior to sequencing Akata lines were removed from G418 selection for 2 weeks and matched to untreated parental lines. LMP-1 transduced (and relevant parental controls) were treated for 24hrs with 10ng/ml Doxycycline to induce LMP-1 expression, prior to RNA extraction. 2 x10^6^ cells were washed twice in PBS before RNA extraction. Cells were homogenised in Trizol (ThermoFisher Scientific) and incubated at room temperature for 5 min. Chloroform was then added to the homogenate (0.2 ml chloroform per ml of Trizol used) and mixed by shaking vigorously for 15 seconds. The sample was allowed to sit at room temperature for 2-3 min, then spun at 12000x g for 15 min at 4°C. The aqueous phase was removed and transfer to new tube. The supernatant was removed and diluted 1:1 with an equal volume of 70% EtOH, the solution was then applied to RNA purification column from a Nucleospin RNA kit (Macherey-Nagel). From there, the manufacturers’ protocol, including on column DNA digest, was followed. RNA was eluted in supplied RNAase free water. RNA concentration was checked by Nanodrop and samples stored at -80°C. RNA was assessed on Tapestation and the bulk RNA sequenced using TruSeq library preparation and the Illumina NextSeq 1000. Resulting reads were aligned to both the human and EBV genomes.

### Bulk RNA extraction & sequencing

RNA was extracted with the RNeasy FFPE Kit (Qiagen 73504) according to the manufacturer’s instructions. Briefly, tissue curls were deparaffinized in xylene and rehydrated. Tissue was digested in buffer PKD with 10 µL of proteinase K and 2 µL of β-mercaptoethanol at 56°C for 15 mins followed by 15 mins at 80°C. Samples were cooled on ice and the digested tissue was pelleted by centrifugation. DNA was digested in DNase Booster Buffer with 10 µL of DNase I at room temperature for 15 mins. The RNA was added to a RNeasy spin column and washed with buffer RPE before eluting in 30 µL of RNase-free water. 5 µL of RNA was removed to determine the quality and quantity of the extracted RNA using the Agilent TapeStation. The remaining 25 µL was used for library prep with the Lexogen QuantSeq 3’ mRNA-Seq Kit FWD for Illumina. QC was evaluated by spiking in 0.5uL of SIRV to allow technical evaluation of library prep and sequencing performance. RNA was sequenced on an Illumina NextSeq 550.

### Bulk & cell line RNA sequencing alignment & analysis

Fastq files were concatenated for each sample replicate. RNA quality control of fastq files was assessed with FASTQC ^43^. The adapter contamination, polyA read through and low-quality tails were trimmed using bbduk, according to Lexogen recommendations of the 3’ assay. Alignment to the GRCh38 Homo sapiens genome was conducted using STAR ^44^, with duplicate reads subsequently removed with PICARD. BAM files were then indexed using SAMtools and counted using HTSEQ-count ^45, 46^. The count files were then imported into R normalised using the median of ratios in DESeq2 ^37^. Gene set enrichment analysis was conducted with the GSEA program.

### THP1 cell culture and treatments

THP1 cells were differentiated into macrophages with 25 ng/mL of PMA, followed by a 48-hour rest period. Subsequently, differentiated THP1 cells (at 70% confluency) were incubated in the supernatant of control, LMP1 transduced and EBV infected SUDHL5 cells for 2 days at 37°C.

### qPCR

RNA was extracted from THP1 cells using the Qiagen RNeasy Plus Mini Kit, followed by cDNA synthesis using the SuperScript VILO cDNA Synthesis Kit (Invitrogen). qPCR was carried out using TaqMan Gene Expression assays for CXCL9 and SPP1, with GAPDH as the endogenous control, and TaqMan Universal PCR Master Mix (Supp. Table S7). Each reaction was performed using 25 ng cDNA as input. No template and no reverse transcriptase controls were included in all experiments.

### Survival Analysis

The RNA sequencing dataset of 1001 DLBCL tumours was obtained from Reddy et al.^47^ Cases with low quality reads were removed from the analysis, leaving 624 DLBCL cases. ABC cases were split into high and low groups for SPP1 using CutOffFinder^48^. Kaplan-Meier analysis was conducted in R using the survival package and visualised with the survminer package.

### Data visualisation and statistics

All data was plotted in R using ggplot2 and ggpubr. Heatmaps were visualised using ComplexHeatmap^49, 50^. The Mann-Whitney U test was used to compare non-Gaussian unpaired data between two groups. For statistical analyses of correlations, the non-parametric Spearman test was used. P values of less than 0.05 (P < 0.05) were considered statistically significant and denoted by the following: *P < 0.05, **P < 0.01, ***P < 0.001 and ****P < 0.0001. Statistics were computed using GraphPad Prism (v10.4.1) or R (v4.0.1). All statistical information can be found in the figures or figure legends. Illustrations were created with BioRender.com.

## Data Availability

Data generated for this study will be made available upon publication or request before publication.

## Acknowledgements

The authors would like to thank Akoya Biosciences for imaging the PhenoCycler-FUSION samples. EBV virus and LMP1 plasmids were kindly provided by Dr. Claire Shannon-Lowe (Birmingham; UK). É.F. is funded by an Irish Research Council Postdoctoral Fellowship (GOIPD/2022/97). THP1 cells were kindly provided by Dr. Jason Bennett (University of Limerick, Ireland). This work was partly funded by the Limerick Digital Cancer Research Centre (LDCRC). Illustrations were created with BioRender.com.

## Author Contributions

É.F. & P.G.M. conceived and designed the study;

G.S.T., A.G.R., S.C.L., & M.K.G. obtained ethical approval for the blood samples;

S.C.L., A.C.D., G.S.T. and M.K.G acquired and processed the blood samples;

S.C.L and A.C.D. conducted the ELISpot experiments;

M.R.P. S.D., L.M., L.L., S.L., & K.H.Y. obtained ethical approval for and acquired the tissue samples;

N.N. stained tissue samples and acquired multiplex imaging data;

A.M.R. extracted tissue RNA and DNA for sequencing;

A.S.H. conducted chromogenic staining;

C.I.L. conducted the IM Tonsil staining and mIF analysis;

C.M.E. conducted the CD20 *in vitro* assays and qPCR analyses;

É.F. conducted all the analysis;

É.F. prepared the figures;

É.F. and P.G.M. wrote the manuscript; M.G.K., G.S.T., & P.G.M. supervised the work.

All the authors contributed to the final version of the manuscript.

